# Association of tamoxifen resistance and lipid reprogramming in breast cancer

**DOI:** 10.1101/276618

**Authors:** Susanne Hultsch, Matti Kankainen, Lassi Paavolainen, Ruusu-Maaria Kovanen, Elina Ikonen, Sara Kangaspeska, Vilja Pietiäinen, Olli Kallioniemi

## Abstract

**Background:** Tamoxifen treatment of estrogen receptor (ER)-positive breast cancer reduces mortality by 31%. However, over half of advanced ER-positive breast cancers are intrinsically resistant to tamoxifen and about 40% will acquire the resistance during the treatment.

**Methods:** In order to explore mechanisms underlying endocrine therapy resistance in breast cancer and to identify new therapeutic opportunities, we created tamoxifen-resistant breast cancer cell lines that represent the luminal A or the luminal B. Gene expression patterns revealed by RNA-sequencing in seven tamoxifen-resistant variants were compared with their isogenic parental cells. We further examined those transcriptomic alterations in a publicly available patient cohort.

**Results:** We show that tamoxifen resistance cannot simply be explained by altered expression of individual genes, common mechanism across all resistant variants, or the appearance of new fusion genes. Instead, the resistant cell lines shared altered gene expression patterns associated with cell cycle, protein modification and metabolism, especially with the cholesterol pathway. In the tamoxifen-resistant T-47D cell variants we observed a striking increase of neutral lipids in lipid droplets as well as an accumulation of free cholesterol in the lysosomes. Tamoxifen-resistant cells were also less prone to lysosomal membrane permeabilization (LMP) and not vulnerable to compounds targeting the lipid metabolism. However, the cells were sensitive to disulfiram, LCS-1, and dasatinib.

**Conclusion:** Altogether, our findings highlight a major role of LMP prevention in tamoxifen resistance, and suggest novel drug vulnerabilities associated with this phenotype.

## Background

Approximately two thirds of breast cancers are estrogen receptor (ER) positive. As the receptor stimulates proliferation of mammary epithelial cells, it is also an important target in anti-hormonal cancer therapy. One of the most prescribed ER antagonists for first line therapy is tamoxifen that has helped millions of women since its discovery 50 years ago [1]. However, *de novo* or acquired drug resistance towards tamoxifen is a notable problem and the later affects approximately 40% of patients receiving tamoxifen [2].

In addition to its intended anti-cancer effects, tamoxifen is known to have both direct and indirect effects on the cellular lipid metabolism. It has been shown to reduce blood cholesterol levels [3] and to be protective against cardiovascular diseases [4]. However, approximately 43% of the patients treated with tamoxifen develop hepatic steatosis, including the accumulation of neutral lipids to lipid droplets in hepatic cells [5]. Tamoxifen can regulate the lipid balance e.g by binding to the microsomal antiestrogen binding sites (AEBS), which are associated with cholesterol metabolism [6]. This mechanism has been linked to control cell growth, differentiation and apoptosis in the presence of reactive oxygen species (ROS) and has been established as another mode by which tamoxifen induces cytotoxicity [7,8].

On the other hand, reprogrammed metabolism is one hallmark of cancer cells [9] and has recently been suggested as a new mode of drug resistance in cancer therapy [10,11]. The metabolic intermediates can supply cancer cells with membrane phospholipids, with energy through the β-oxidation pathway or with pro-tumorigenic lipid-signaling molecules such as lysophosphatidic acid [12]. Some studies even suggested a role of cholesterol metabolism in tamoxifen resistance [13] and Borgquist et al. showed an improved clinical outcome for ER-positive breast cancer patients receiving cholesterol-lowering medication during their adjuvant endocrine therapy [14].

In the present study, we have delineated mechanisms underlying tamoxifen resistance by extending our drug screening and exome sequencing analyses on tamoxifen-resistant cell lines [15] by performing RNA-sequencing on the tamoxifen-resistant and their isogenic, tamoxifen-sensitive parental cell lines, and searched for genes and pathways that may be involved in the acquired resistance. Differential gene expression and pathway analysis confirmed that endocrine resistance is not triggered by one common mechanism but involves several functional pathways, depending on the cell type [15]. Through the integration with public data [16], the usefulness of our breast cancer cell line model was assessed and the relevance of the identified transcriptomic alterations verified in this patient cohort. By focusing on the isogenic T-47D cell variants, we showed that genes in the cholesterol pathway were differentially expressed between tamoxifen-sensitive and tamoxifen-resistant cells. These results were supported by a striking accumulation of lysosomal cholesterol upon tamoxifen treatment, increased neutral lipid amounts after the development of resistance. Markedly, tamoxifen treated cells were also found to be less prone to lysosomal membrane permeabilization, suggesting that altered lysosomal integrity may confer resistance to tamoxifen. Finally, using high-content phenotypic drug sensitivity profiling of 33 drugs targeting lipid metabolism as well as inducing lysosomal membrane leakage, we identified drug candidates for overwriting the tamoxifen resistance and potentially being beneficial for breast cancer patients unresponsive to tamoxifen.

## Methods

### Cell culture

The development, characterization and culturing of the tamoxifen-resistant and their isogenic parental cell lines has been published previously [15]. If not otherwise stated, the parental cell lines (luminal A: MCF-7, T-47D, ZR-75-1, luminal B: BT-474) [17] were cultured without tamoxifen and the tamoxifen-resistant cell lines, marked with Tam throughout this study (MCF-7 Tam1, T-47D Tam1 & Tam2, ZR-75-1 Tam1 & Tam2, BT-474 Tam1 & Tam2) were supplemented with 1 μM 4-OH-tamoxifen (Sigma).

### RNA-sequencing and data analysis of cell line data

RNA isolation, library preparation, sequencing, and data-analysis were done as explained in Kumar A et al. [18]. Briefly, total RNA was isolated using miRNeasy kit (Qiagen) and its quality was controlled by using the Agilent Bioanalyzer with the RNApico chip (Agilent). Qubit RNA-kit (Life Technologies) was used to quantitate RNA amount per sample. The strand-specific paired-end RNA-sequencing library was prepared then with ScriptSeq™ Complete kit for human/mouse/rat (Illumina). The library preparation included the ribodepletion of rRNA from 1 μm of total RNA and generation of double stranded cDNA by revers transcription with random hexamers for generation of cDNA. SPRI beads (Agencourt AMPure XP) were used to purify the libraries and to remove fragments less than 200 bp in length. The mean fragment size ranged from 300 to 400 nucleotides. The library quality was evaluated on the High Sensitivity chip by Agilent Bioanalyzer (Agilent). The paired-end sequencing was performed using the Illumina HiSeq 2000 (Illumina) instrument according to the manufacturer’s instructions.

RNA-sequencing data analysis was performed as described in detail in Kumar et al. [18] and included preprocessing of read data, gap-aware alignment of the read data to the human reference genome (Ensembl GRCh38) with the guidance of the EnsEMBL reference gene models (EnsEMBL v80), read summarization against EnsEMBL v80 gene features, and identification of fusion genes. Fusion genes were detected using FusionCatcher, which was applied to raw, un-processed read files with default parameters [19]. The raw and processed sequencing data have been deposited in the GEO database [GEO: GSE111151].

### Integration of public and cell-line transcriptomic data and differential expression analysis

Public patient transcriptome data [16] was downloaded from GEO database [20] and analyzed as cell line sequencing data, but using the EnsEMBL v82 gene features in all steps. Alignment files were combined with alignment files from the cell-line samples and count estimates were generated against EnsEMBL v82 gene features using subreadR [21]. Count data was then assigned to EnsEMBL information using biomaRt [22], normalized using the trimmed mean of M-values (TMM) method [23], converted to CPM (counts per million) estimates using edgeR [24], corrected for the batch effect associated with the study origin using limma [25], and filtered for lowly expressed features showing log2 expression ≤ 1 in over half of samples. Default parameters were used. In the lack of biological replicates, differentially expressed genes were identified as those with a log2 ratio of >= |10| and CPM difference of >= |10| against its matched parental cell line. Further, we used Enricher [26,27] with the list of differentially expressed genes to determine pathways that were involved in the development of tamoxifen resistance for each parental vs. resistant cell line comparison. Pathways with an adjusted p-value less than 0.001 (1E-3) were accepted as significantly altered. Log2 ratios of differentially expressed genes of the cell lines were visualized using heat maps. In the heat map analysis, genes and samples were ordered using unsupervised complete linkage clustering with Euclidean distance measure [15]. Patient and cell line data was visualized using principle component analysis and heat maps as described above using biomaRt to extract cholesterol genes under the reactome.ID R-HSA-191273.

### Measurement of triglycerides and cholesterol esters

Parental and resistant T-47D cells were plated on 6-well plates and grown in their default media +/− 1 μM 4-OH-tamoxifen for 72 h in triplicates. Cells were scraped in 0.2 N NaOH for lipids extraction as previously described [28]. Cell lysates were also examined for protein amount. Free cholesterol, cholesterol esters and triglycerides were resolved on TLC plates using hexane/diethyl ether/acetic acid (80:20:1) as the mobile phase prior to the visualization of lipids by charring. The lipid bands were quantified by ImageJ [29] from scanned plates and the lipid amounts were determined based on the standard curves for triglycerides, cholesterol esters and free cholesterol.

### Immunofluorescence staining and Western blotting

Parental and resistant T-47D cells were seeded on coverslips and grown in the default media +/− tamoxifen for 72 h. Cells were fixed with 4 % paraformaldehyde for 20 minutes at room temperature, and permeabilized with 0.3 % Triton X-100 for 5 minutes, followed by 30 min blocking with 3 % BSA-PBS at 37 °C. The primary and secondary antibodies were diluted in 1 % BSA-PBS and consecutively incubated for 60 and 30 min at 37 °C. For detection of free cholesterol, cells were stained with 0.05 % filipin (Sigma) in 10 % FBS-PBS blocking solution for 30 minutes at 37 °C. Antibodies were prepared in 5 % FBS-PBS and incubated as described above. Nuclei were stained with DRAQ5 (Biostatus). For detecting lipid droplets, LipidTOXGreen neutral lipid stain (ThermoFisher Scientific) was diluted 1:200 in PBS to stain freshly fixed coverslips following the manufactures protocol and detecting the nuclei with Hoechst. Stained coverslips were mounted with Prolong Gold anti-fade reagent (Invitrogen) and imaged with a Nikon 90i microscope (Nikon). For Western blotting, cells were grown on 10 cm dishes, and lysed in Laemmli buffer. Immunoblotting was performed as previously described [30] using the Odyssey Blocking Buffer (Licor) for blocking and the antibody dilutions. Information about the antibodies and their dilution used for immunofluorescence as well as Western blotting is available at Additional File 1.

### Lysosomal membrane permeabilization (LMP) assay

In order to measure the integrity of lysosomal membranes, we performed the detection of damaged lysosomes by galectin-1 and −3 translocation according to the previously published protocol [31]. We first established that galectin-3 was in our cell lines a more reliable marker to detect its translocalization to the lysosomes compared to galectin-1. We then seeded 2000 cells/well of T-47D, T-47D Tam1 and Tam2 on PE Cell-Carrier 384-well plate +/− 1 μM 4-OH-tamoxifen. After 72 h incubation, 1 mM LLMOe was added as LMP induction control and incubated for 1 h. Cells were then fixed with 4 % PFA and stained with galectin-3 detected with Alexa 568 (461nm), ceruloplasmin as cell segmentation marker detected with Alexa 488 (488 nm, Additional file1) and Hoechst (405nm) for detection of the nuclei. 16 fields-of-view were acquired with the two sCMOS cameras (2160×2160 pixels) containing Opera Phenix HCS system (PerkinElmer) in a widefield mode with the 40x water immersion objective (NA 1.1). Exposure time and laser power were kept constant for each individual staining across different cell lines and conditions. We utilized the Columbus Image Data Storage and Analysis System (PerkinElmer) to analyze the multi-channel images. First, the images were preprocessed to correct non-uniform illumination. Next, individual nuclei were segmented from the Hoechst channel. The minimum area of a nucleus was set to 30 μm^2^ to remove the detection of debris in the image background. Starting from the detected nuclei, the segmented regions were propagated to cover cell cytoplasm stained with ceruloplasmin. The cells that touched the image border were discarded. Finally, spot detection was used to segment galectin-3 stained spots. The maximum radius of the spots was set to 1 μm. We defined cells with more than 1 spot as galectin-3 positive to exclude false positive detection. Further, we calculated the percentage of galectin-3 positive cells and the average spots per cell.

### Drug sensitivity and resistance testing (DSRT) and high-content phenotypic drug profiling

33 compounds that target lipid and cholesterol metabolism, or induce LMP, were selected for the DSRT by literature and vendor research (Additional file 2). As in the previous DSRT screens [15,32], the dissolved drugs were transferred in five different concentrations covering a 10 000-fold concentration range into 384-well plates in duplicates mirrored after column 12. 1500 cells of T-47D parental, Tam1, and Tam2 cells were then seeded into the wells in normal growth media on columns 1-12 of each plate and in media supplemented with 1 μM 4-OH-tamoxifen on columns 13-24. The cells were then incubated for 72 h at 37 °C and cell viability was measured by CellTiter-Glo (CTG) Cell Viability Assay (Promega) with the PHERAstar plate reader. Data was normalized to negative (DMSO only) as well as positive (100 μmol/l benzethonium chloride) controls, logistic dose response curves fitted using the Marquardt-Levenberg algorithm, and Drug Sensitivity Score (DSS) calculated as described previously [33], and implemented in the in-house bioinformatics analysis pipeline Breeze.

For the high-content phenotypic drug profiling, plates were fixed with 4 % PFA-PBS after incubation with drugs for 72 h, and stained with LipidTOXGreen neutral lipid stain (ThermoFisher Scientific) diluted 1:200 in PBS and Hoechst for nuclei detection. 25 fields-of-view per well were acquired with the PE Opera Phenix HCS system (PerkinElmer) in a confocal mode with the 40x water immersion objective (NA 1.1). Exposure time and laser power were kept constant for each individual staining across different cell lines and conditions. Images were analyzed using a custom pipeline to measure LipidTOXGreen signal and image-based DSS based on cell counts. Images were preprocessed by applying flatfield correction using CIDRE method [34], and then stitching corrected 25 fields-of-view images to a single image of a well for each channel. The stitching was done to enable the analysis of cells crossing internal borders of images. The stitching produced images of approximately 10400^2^ pixels in size which were resampled to 5200^2^ pixels to reduce computational capacity needed for image analysis. Images were analyzed with CellProfiler 2.2.0 [35]. First, nuclei were segmented from the Hoechst channel using Otsu thresholding followed by separation of touching nuclei with watershed transform on distance transformed image. LipidTOXGreen channel was segmented using adaptive Otsu thresholding and propagation outwards from individual nuclei. The LipidTOXGreen signal was measured in segmented regions for each individual cell. Logistic dose response curves were fitted to cell counts to calculate image-based DSS in Breeze.

### Statistical analysis

For all experiment that were at least done in triplicates the values were expressed as mean ± SD. One way ANOVA was performed on the mean of each measurement followed by Tukey test to enable multiple comparisons between groups. Statistically significance was accepted as p<0.05. All comparisons of the measurement of triglycerides, free cholesterol and cholesterol esters, LMP assay, and LipidToxGreen staining can be found in Additional file 3.

## Results

### Each tamoxifen-resistant cell line develops its individual gene expression and fusion gene profile

RNA-sequencing was performed to detect differentially expressed genes and pathways between parental and resistant cell lines. Between 85 and 181 million filtered reads were obtained per sample (Additional file 4), providing means to obtain expression estimates between 33,600 and 37,000 genes per sample. We determined differential gene expression as log2 change of >|1|, and the difference of gene expression > |10| CPM between the resistant clone and its isogenic parental cell line (see Methods). Using this filtering, we identified >1200 differentially expressed genes in MCF-7-and T-47D-derived cells, and <400 genes in BT-474-and ZR-75-1-derived cells. On average 59 % of differentially expressed genes were upregulated in the majority of resistant cell lines (Table 1). Interestingly, only about 35 % of altered genes in T-47D as well as BT-474, and only 24 % in ZR-75-1 were shared between the resistant cells derived from the same parental cell line (Additional figure 1A). Additionally, no common differentially expressed genes were identified, highlighting that each of the cell lines had developed resistance through a distinct molecular pattern (Figure 1A and B). Nevertheless, common genes (*SERPINA1, PLXDC2, NAV1, DOCK1O* and *LRP1*) with altered expression were detected in the luminal A cell lines (Additional figure 1B). No shared fusion genes across the cell lines were detected apart from the read-through fusions *ABCC11-ABCC12* and *TRIM3-HPX* (Additional file 5).

**Figure 1:**
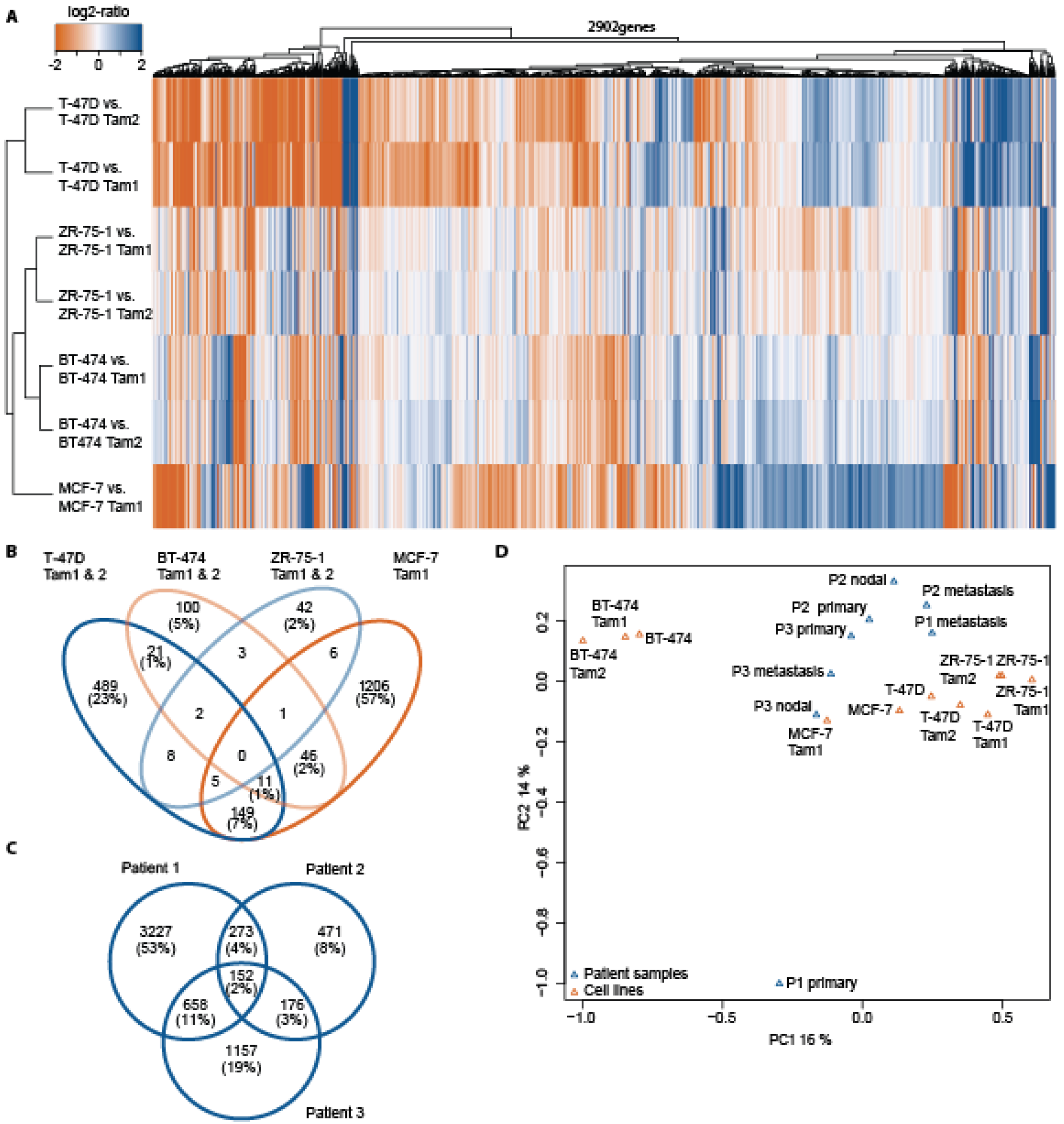
Tamoxifen-resistant cell lines display distinct expression changes and share similarities with patient cases. **A** Hierarchical clustering and heat map visualization of each parental/resistant cell line pair. Orange (negative log2-ratio) represents increase and blue (positive log2-ratio) decrease in expression in the resistant cell lines. Only protein coding genes with log2 ratio >|1| and the difference of gene expression > |10| CPM in at least one of the comparisons are displayed. Log2 ratios of >= |2| are displayed in the same color. Tamoxifen-resistant clones (**B**) and patient samples (**C**) differ in their expression changes. Parental is compared with resistant cell line and primary tumor with metastatic tumor from the same patient, respectively. Venn diagrams show overlap in numbers and percentage of genes that are differentially expressed. **D** PCA plot indicates that patients share expression patterns with the luminal A cells.

**Table 1.** Differentially expressed genes

| Number of | T-47D<br>Tam1 | T-47D<br>Tam2 | BT-474<br>Tam1 | BT-474<br>Tam2 | MCF-7<br>Tam1 | ZR-75-1<br>Tam1 | ZR-75-1<br>Tam2 |
| --- | --- | --- | --- | --- | --- | --- | --- |
| differentially<br>expressed genes | 1425 | 1239 | 303 | 360 | 1424 | 193 | 158 |
| upregulated genes | 941 | 823 | 195 | 183 | 640 | 104 | 68 |
| downregulated<br>genes | 484 | 416 | 108 | 177 | 784 | 89 | 90 |

### Tamoxifen-resistant cell lines resemble tamoxifen-treated patient cases

To compare the tamoxifen-resistant cell line models with breast cancer patient samples, we reanalyzed the McBryan et al. RNA-sequencing data [16]. Following the analysis with our RNA-sequencing pipeline and correction for batches (patient samples and cell lines), we determined differentially expressed genes between primary and metastatic tumors and integrated the data with the transcriptome data from our cell lines. The results were in line with McBryan’s et al., and indicated that only about 2.5 % of differentially expressed genes were shared between all the patients (Figure1 C). Interestingly, we found that patient-specific expression profiles clustered together with the expression of the luminal A cells (Figure 1D). Furthermore, *SERPINA1*, which was differentially expressed in the tamoxifen-resistant luminal A cell lines (being upregulated in MCF-7 Tam1 and downregulated in T-47D Tam1, T-47D Tam2, ZR-75-1 Tam1 and ZR-75-1 Tam2), also changed its expression in the patient samples (being up-regulated in all three metastatic tumors). *CP* (Ceruloplasmin), a gene that was highly upregulated in the resistant T-47D cell lines (240-290 fold-increase) as well as MCF-7 Tam1 (26 fold-increase), was also found to be overexpressed in all the 3 metastatic patient samples ranging from 12-fold increase (patient 2) to 50-57 fold increase (patient 1 and 3, respectively, Additional file 6).

### Triglycerides and cholesterol esters are increased in the resistant T-47D cell lines

To reveal pathways associated with tamoxifen resistance, we analyzed the differentially expressed genes with Enrichr [26,27]. Based on Enrichr′s Reactome 2016 analysis with an adjusted p-value below 0.001, we observed multiple enriched pathways in different resistant cell lines (Table 2 and Additional File 7). The most striking differences were found in the T-47D Tam1 and Tam2 cells, which displayed changes in metabolism associated genes, especially those involved in cholesterol and related lipid metabolism (Table 2, Figure 2A).

**Figure 2:**
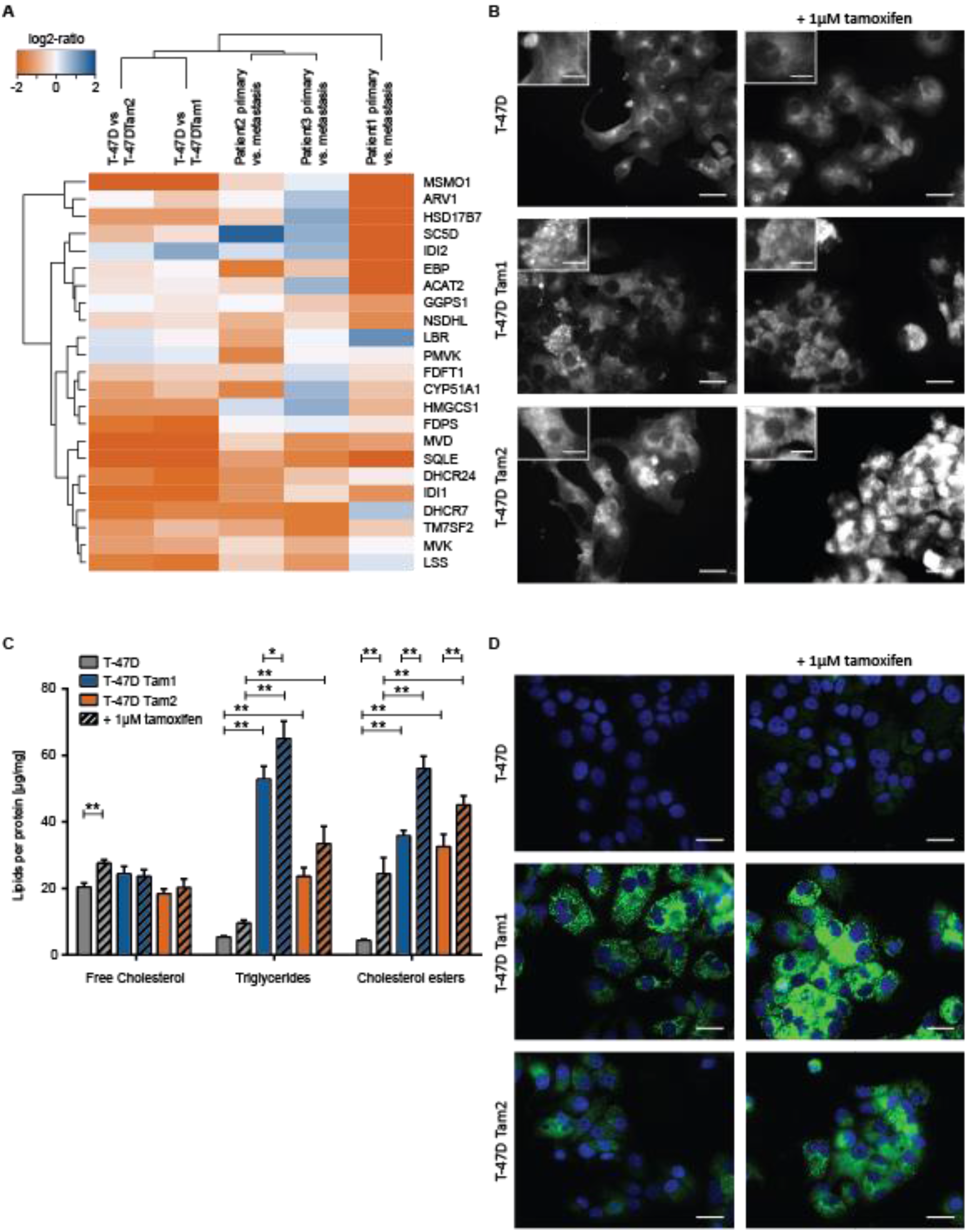
Genes of cholesterol pathway and related lipids are upregulated in the tamoxifen-resistant T-47D cell lines. **A** Hierarchical clustering and heat map visualization of each parental/resistant cell line pair and patient primary/metastatic tumor. Orange (negative log2-ratio) represents increase and blue (positive log2-ratio) decrease in expression in the resistant cell lines/metastatic tumor. Log2 ratios of >= 121 are displayed in the same color. **B** Filipin staining reveals an increase in intracellular amounts of free cholesterol in tamoxifen-resistant T-47D cells +/− 1μM 4-OH-tamoxifen. **C** Quantification of lipid content in cells grown +/− 1μM 4-OH-tamoxifen reveals an increase in cholesterol esters and triglycerides in tamoxifen-resistant cells (depicted as colored bars). Only significant differences (p-value <0.05) between the same clone as well as of the comparison between resistant and tamoxifen-resistant cells in the same treatment conditions are indicated all other comparisons can be found in Additional File 3. **D** LipidToxGreen staining of neutral lipids (green) demonstrates that accumulation of neutral lipids into lipid droplets in tamoxifen-resistant cells. The nuclei (blue) were stained with Hoechst.

**Table 2.**
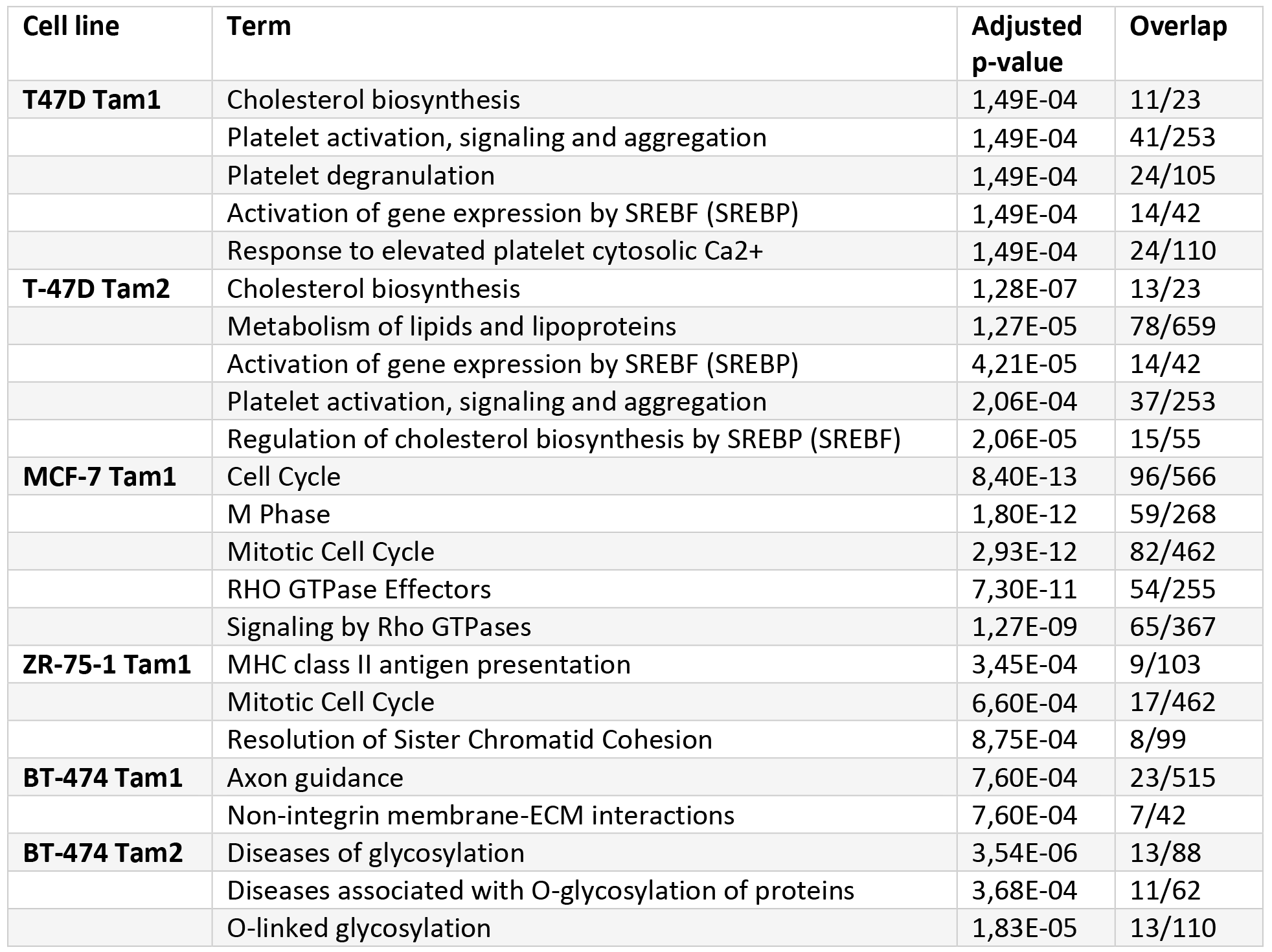
TOP5 Pathways with adjusted p-value below 0,001 obtained from Enricher using the Reactome 2016 pathway. Resistant cell lines are compared with the isogenic parental control cells.

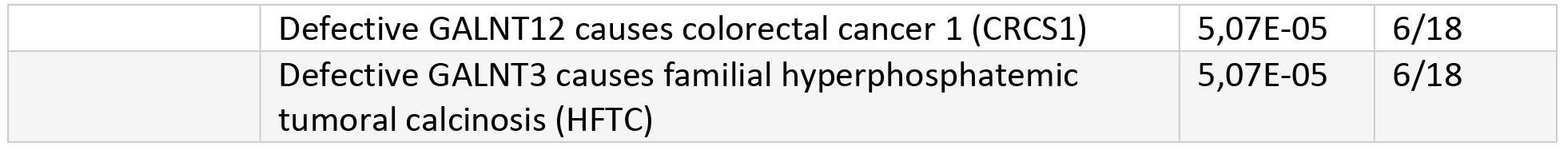

In addition, we observed an upregulation of genes involved in cholesterol biosynthesis in all three metastatic patient samples from McBryan et al. study (Figure 2A). Therefore, we focused our studies on these pathways within the T-47D cell lines (Table 2 and Additional File 7). To investigate whether deregulation of genes involved in cholesterol biosynthesis could affect the cellular cholesterol balance, we stained cellular free cholesterol with filipin, a fluorescent cholesterol-binding compound. Notably, we observed increased intracellular amounts of free cholesterol in the resistant cells, displaying a cumulus cloud-like staining pattern (Figure 2B). To quantify the presence of major cellular lipid species e.g. cholesterol, cholesterol esters, and triglycerides, their amounts were further determined with thin layer chromatography. The total cellular free cholesterol remained unchanged, suggesting that only the distribution of free cholesterol was altered in the resistant cells. However, we observed an increase in neutral lipids (cholesterol esters and triglycerides) upon tamoxifen treatment. The increase in triglycerides was significantly high (4 to 7 fold-increase) in resistant cells compared to parental cells (Figure 2C). To visualize the changes in neutral lipid amounts as well as their intracellular distribution, we stained the cells with LipidToxGreen, a fluorescent dye binding specifically to neutral lipids. The analysis indicated that most of the neutral lipids accumulated in enlarged lipid droplets, which fill the cytoplasm (Figure 2D). Moreover, RNA sequencing results implicated that the expression of Peroxisome Proliferator-Activated Receptor gamma (*PPARG*), which is known to regulate several lipid droplet proteins, was upregulated in resistant cells. In addition, the ATP Binding Cassette Subfamily A Member 1 (*ABCA1*), which functions as a cholesterol efflux pump was downregulated (Additional file 6).

### Tamoxifen-resistant cells show altered morphology of lysosomes, have altered processing of Cathepsin D, and are less susceptible to lysosomal membrane permeabilization

To localize the accumulated free cholesterol, we co-stained free cholesterol (filipin) with antibodies detecting the lysosomal-associated-membrane-proteins 1 and 2 (Lamp1 and Lamp2). Based on this analysis, we observed that most of the free cholesterol accumulated into lysosomes. We also discovered an increase in the amount and size of lysosomes as well as divergences in shape compared to their typical round form seen in the parental cell lines (Figure 3A & B). As the tamoxifen-resistant cells displayed a prominent phenotype with free cholesterol accumulation to structurally disturbed lysosomes, we studied the amounts of cathepsin D and its lysosomal maturation. Cathepsins are lysosomal proteins that help to maintain the homeostasis of cell metabolism and are involved in apoptotic signaling as well as in lysosomal membrane permeabilization. Furthermore, the expression of cathepsin D is known to be regulated by estrogen [36]. As expected, we observed a decrease of mature cathepsin D (28 kDa) under tamoxifen treatment in parental and resistant cell lines, suggesting that tamoxifen can regulate the expression and/or processing of cathepsin D [37]. Whilst addition of tamoxifen also caused an upregulation of the precursors of cathepsin D in the parental cell line, such an increase was not obvious in the resistant cell lines (Figure 3E), suggesting that the maturation of cathepsin D in the lysosomes may be affected.

**Figure 3:**
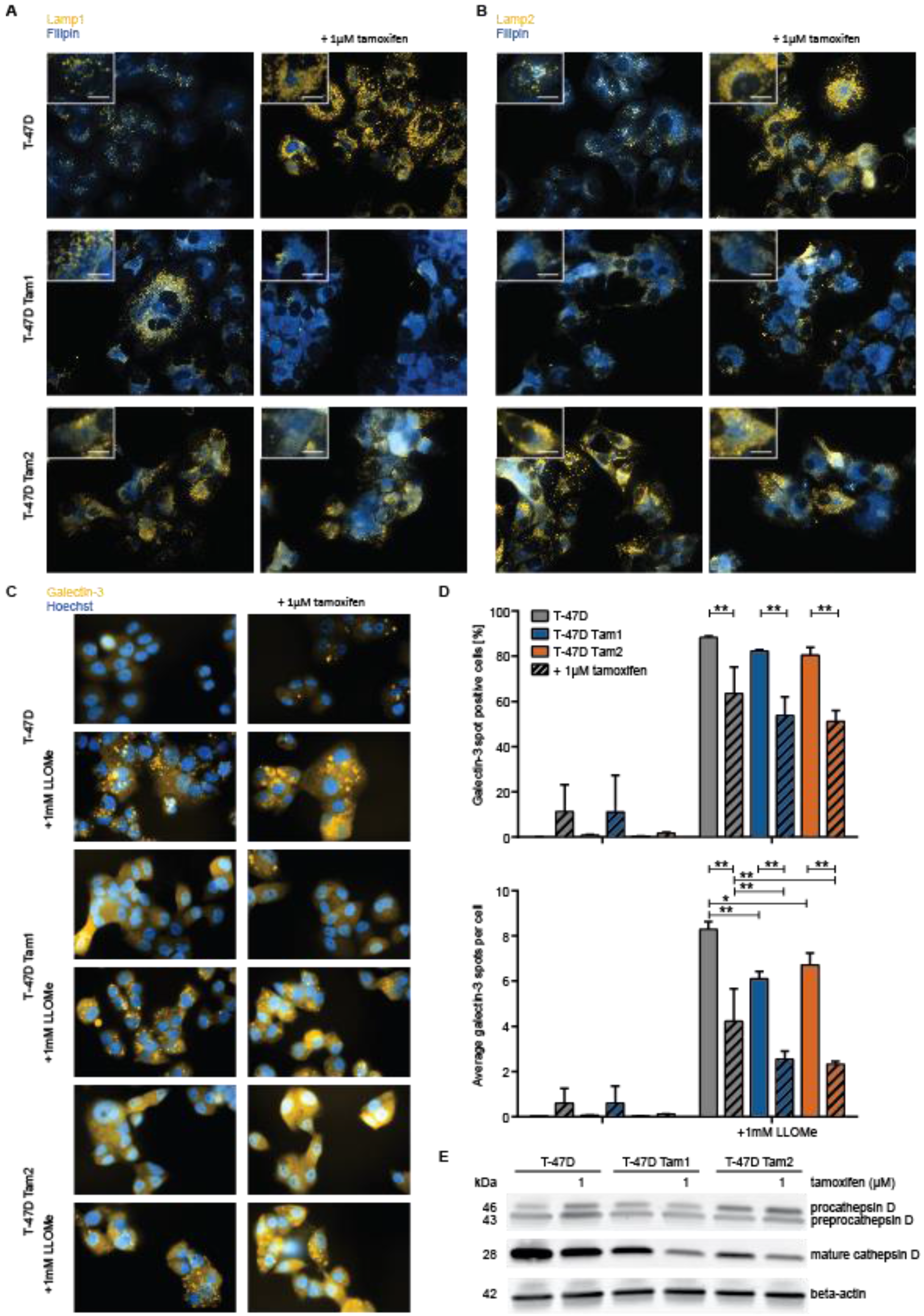
Free cholesterol accumulates in lysosomes in the resistant cells. Intracellular accumulation of free cholesterol (blue) accumulates in lysosomes (orange) stained with two lysosomal markers, Lamp1 (**A**) and Lamp2 (**B**), detecting the lysosomal-associated-membrane-proteins 1 and 2 +/− 1μM 4-OH-tamoxifen. Tamoxifen-resistant cells are less sensitive to lysosomal membrane premeabilization detected with galectin-3 (orange) translocation (**C**, images were differently enhanced for visualization purposes) and measurement of galectin-3 positive cells (**D** upper graph) as well as number of galectin-spots per cell (**D** lower image). Galectin-3 measurements were done on the raw image. Only significant differences (p-value <0.05) between the same clone as well as of the comparison between resistant and tamoxifen-resistant cells in the same treatment conditions are indicated all other comparisons can be found in Additional File 3. **E** Mature cathepsin D is downregulated in tamoxifen resistant cells.

As lysosomal integrity, with cholesterol content of lysosomal membranes and cathepsin D among its regulators, plays an important role in the induction of cell death [38], we monitored the translocation of galectin-3 to detect LMP. Given that galectin-3 translocation to the lysosomes was not detected in the parental cells when grown without tamoxifen, and that only very few tamoxifen-treated cells showed galectin-3 spots (Figure 3C & D), lysosomes were most likely undamaged and functional in all cells. Further, by inducing LMP with 1 mM LLOMe we were able to observe that tamoxifen treated resistant cells were less susceptible to LMP compared to the parental cell line, having only 51-53% of cells with galectin-3 spots under tamoxifen treatment and significantly less galectin-3 spots per cell (Figure 3C & D). This suggests that circumvention of LMP in the resistant cells leads not only to tamoxifen resistance but may also decrease their sensitivity to other drugs.

### Drug testing of tamoxifen resistant cells reveals sensitivity to dasatinib, disulfiram and LCS-1

Guided by our RNA-sequencing results we selected 33 drugs, known to affect the genes or pathways involved in lysosomal alterations and lipid metabolism as well as some drugs identified in our previous screen (Additional file 2, [15]). As readouts for the DSRT, we applied both enzymatic cell viability measurement (CTG) as well as a phenotypic image-based analysis using LipidToxGreen to observe neutral lipids in lipid droplets together with Hoechst to detect nuclei.

The cell viability measurement revealed drugs that reduced ATP levels in tamoxifen-resistant cells similarly to the control cells, independently of their lipid accumulation phenotype (Figure 4A, Additional file 8).

Dasatinib, a dual Abl/Src inhibitor was more effective in killing the tamoxifen resistant cell lines compared with the parental cells in agreement with our previous results [15]. Tamoxifen-resistant cells were more sensitive to microtubule depolymerizing drugs, such as vincristine and vinorelbine, when measured by ATP amounts. Interestingly, the T-47D Tam2 cells were especially sensitive to vinorelbine induced cytotoxicity (Figure 4A, Additional file 8). The mitotic inhibitors paclitaxel and docetaxel (microtubule stabilizers) were less effective in the T-47D Tam1 cells. (Figure 4A, Additional file 8).

**Figure 4:**
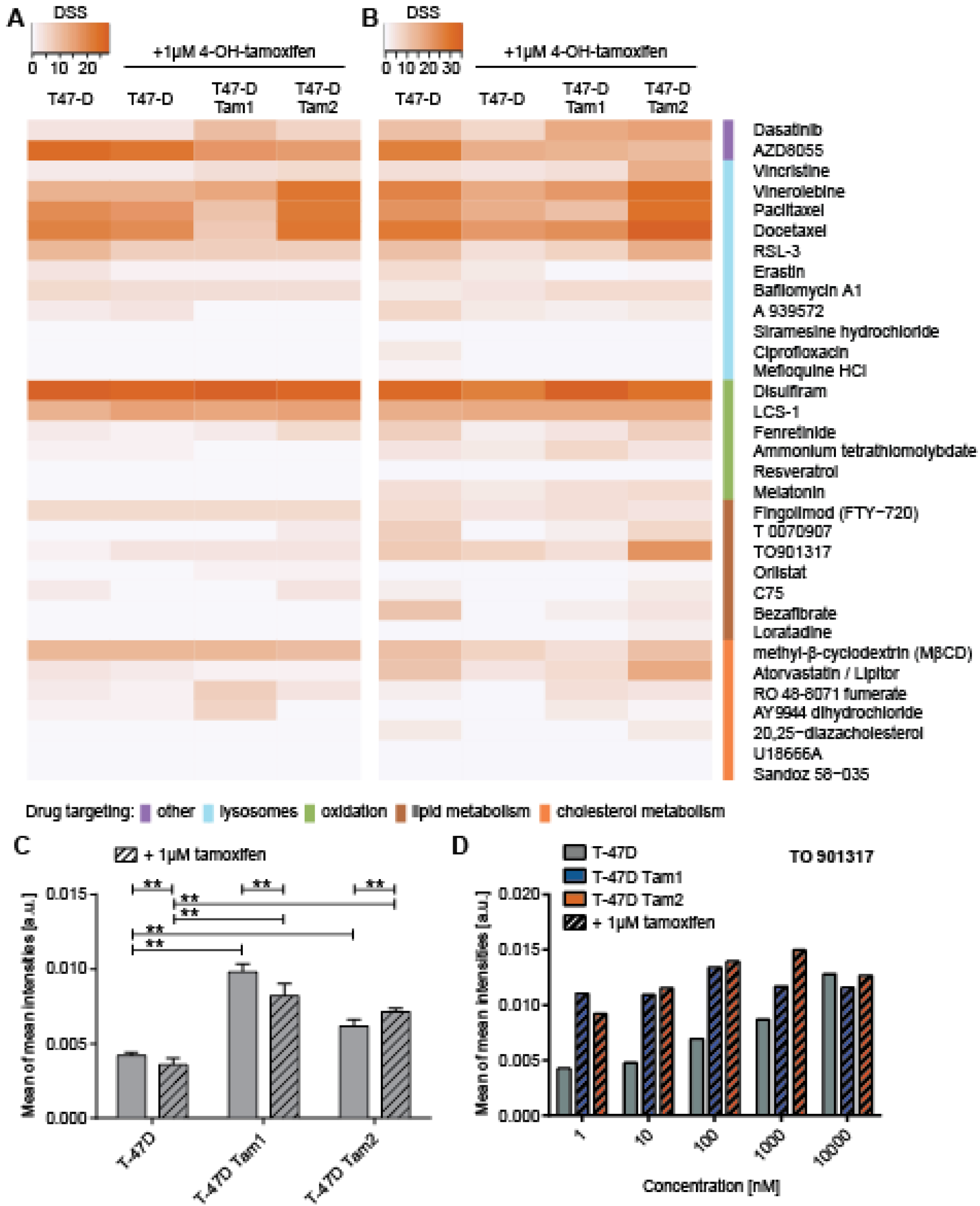
Drug sensitivity and resistance testing of tamoxifen resistant cells. DSS scores of **A** CTG-measurement and **B** cell count are displayed as heatmap with dark orange showing the most effective drugs. **C** Quantification of LipidToxGreen staining in cells grown +/− 1μM 4-OH-tamoxifen reveals an increase in the mean intensities of the staining in tamoxifen resistant cell lines. Only significant differences (p-value <0.05) between the same clone as well as of the comparison between resistant and tamoxifen-resistant cells in the same treatment conditions are indicated all other comparisons can be found in Additional File 3. **D** TO901317 increases the LipidToxGreen staining mainly in the parental cell lines

The most effective drug against all the T-47D clones was disulfiram, a specific inhibitor of aldehyde-dehydrogenase (ALDH1). All the clones responded to AZD8055, a dual mTOR inhibitor, although T-47D Tam1 and Tam2 showed reduced sensitivity. Atorvastatin, which inhibits HMGCoA reductase, did not affect the CTG DSS levels (Figure 4A) but was able to reduce the cell count in the parental and even more in the T-47D Tam2 cells (Figure 4B). The SOD-1 inhibitor LCS-1 was effectively killing both parental and resistant T-47D clones, and RSL-3, a ferroptosis activator due to inhibition of glutathione peroxidase 4, induced cell death in all the cell lines, with somewhat reduced response in T-47D Tam1 (Figure 4B, Additional file 8).

To see whether any of the compounds are able to revert the lipid phenotype prior to reducing the cell viability (ATP-measurement) or induction of cell death (cell count), we specifically monitored the changes in neutral lipids by quantifying the average LipidToxGreen intensity per well (Additional file7). The measured significant increase in the intensity was within the 2-fold range, and we were able to confirm the trends from the biochemical screen where we observed a neutral lipid accumulation in the resistant cell lines (Figure 2C, Figure 4C). Whereas most of the drugs had minor effects on the LipidToxGreen intensity (Additional file 9), the LXR-agonist TO901317 increased the lipid phenotype most strikingly in the parental cells, with less increase particularly in T-47D Tam1 cells (Figure 4D). In addition, methyl-β-cyclodextrin, a membrane cholesterol-depleting agent, caused a lipid droplet accumulation phenotype, mostly in the parental cell line prior to cell killing in the highest concentration (Additional file 9).

## Discussion

In this study, we utilized RNA-sequencing and pathway analysis to understand the underlying tamoxifen resistance and identify resistance-specific drug vulnerabilities. We revealed the involvement of lipid metabolism in tamoxifen resistance as well as pointed out potential therapeutic ways to target these pathways.

Gene expression analysis on tamoxifen-resistant cells reinforced our previous finding on breast cancer cells using a variety of molecular pathways as they acquire tamoxifen resistance [15]. The difference in gene expression was reflected in the scale and scope of differentially expressed genes, and in the lack of shared genes across all the cell lines (Figure 1). In agreement with this finding, the only study that has performed sequential tumor transcriptome analysis on patients developing endocrine resistance, also identified less than 3% of differentially expressed genes across patients (Figure 1C [16]). Despite the overall transcriptome profiles being distinct across the resistant cell lines, we were able to identify five genes that were concordantly differentially expressed in the luminal A subtype resistant cells (Additional figure 1B). Of these, *SERPINA1,* encoding for a serine protease inhibitor primarily targeting elastase, is known to bind ER in a 17β-estradiol (E2) - independent manner, which leads to an increase in its expression [39]. Therefore the observed expression changes could be due to the down-and upregulation of ER in these cell lines [15]. Interestingly, in all three metastatic samples from the McBryan *et al.* study, we observed an increase in *SERPINA1,* which is accompanied by a slight increase of *ESR1* transcription (Additional file 6). Pathway analysis of the differentially expressed genes identified several paths involved in acquired tamoxifen resistance (Table 2, Figure 2A).

In this study, we investigated the tamoxifen-induced changes observed in lipid metabolism, which occurred in the T-47D tamoxifen-resistant cell lines (Table 2, Figure 2). We also made the equivalent finding in a patient′s metastatic tissue (Figure 2A). As the metastasis was found in the liver [16], the observed lipid metabolism pathway profiles have to be interpreted with caution. Nevertheless, our findings suggest that the lipid phenotypes could already develop in the breast cancer cells [40] and is not solely induced by the liver environment.

Further, our studies with the T-47D tamoxifen-resistant cell lines show an increase of free cholesterol into strikingly enlarged lysosomes (Figure 2B, Figure 3A & B, [41]). It has been shown that accumulation of cholesterol, an increase in Lamp1 and Lamp2 as well as downregulation of cathepsins prevents lysosomal membrane permeabilization [42–45], a process which leads to different forms of cell death such as apoptosis, necroptosis, necrosis and ferroptosis [38]. Indeed, our data on the resistant cells shows an increase in cholesterol, Lamp1 and Lamp2, as well as a decrease in cathepsin D (Figure 2B, Figure 3A, B and E [37]). A short-term tamoxifen treatment diminished directly the LLOMe-induced LMP. The T-47D Tam1 and Tam2 were even more resistant towards LMP (Figure 3C and D), showing that tamoxifen can hinder it, and in acquired resistance, this phenomenon is even more prominent. Thus, impeded lysosomal membrane permeabilization may additionally enhance the co-resistance to other cancer drugs during acquired tamoxifen resistance.

Reducing the reactive oxygen species (ROS) is another mechanism by which cells avoid lysosomal induced cell death [44]. We speculate that resistant T-47D cells are able to reduce oxidative stress by upregulation of *SOD1* (Additional file 6) and may therefore be less sensitive to lysosomal cell death. This hypothesis is further supported by the fact that the resistant cells were highly sensitive to the SOD1 inhibitor LCS-1. The capability of erastin to activate ferroptosis is instead inhibited by antioxidants, and it was more effective in parental than in resistant cells. The ferroptosis activator RSL-3, which inhibits the glutathione peroxidase 4, an enzyme that protects from oxygen damage, induced cell death in all the cell lines (Figure 4 and Additional file 8). This further supports the assumption that the T-47D cells are able to reduce oxidative stress and are therefore less sensitive to lysosomal cell death.

Disulfiram, which targets ALDH1 to increase oxidative stress, was highly effective in both parental and tamoxifen-resistant T-47D cell clones (Figure 4 and Additional file 8). The effectiveness of disulfiram is currently investigated in metastatic breast cancer in a phase II clinical trial [46]. *ALDH2*, another target of disulfiram, is upregulated in T-47D Tam1 but not in Tam2 (Additional file 6). High levels of ALDH1 have been shown to predict resistance in women treated with tamoxifen [47], but as *ALDH1A1* is expressed at very low levels in the T-47D cell lines (Additional file 6), we assume that the sensitivity to disulfiram could be due to its capability to disable antioxidation mechanisms of the cells [48].

A significant increase in triglycerides, stored in large lipid droplets (LDs) was observed in tamoxifen-resistant cells (Figure 2C, D and Figure 4C). Free fatty acids are enzymatically converted to triacylglycerol, and then incorporated into LDs. Packaging of excess lipids into LDs could be seen as an adaptive response to fulfilling energy supply without hindering mitochondrial or cellular redox status and keeping the concentration of lipotoxic intermediates low [49]. Accordingly, high LDs and stored cholesterol esters in tumors are considered as hallmarks of aggressive cancer [50]. LD-rich cancer cells have also been shown to be more resistant to chemotherapy [51]. We found over 3-fold upregulation of stearoyl-CoA desaturase (SCD), encoding for a rate-limiting enzyme in the biosynthesis of monounsaturated fatty acids, in the tamoxifen-resistant T-47Ds (Additional file 6). Whether it alone is able to induce the increase in TGs, remains to be investigated. In line with this speculation, *SCD* overexpression has been observed, in different cell types as well as in tamoxifen-induced hepatocyte steatosis, to significantly increase the rate of triglyceride synthesis [52]. The compounds directly affecting lipid metabolism, such as C75, Bezafibrate, T 0070907, TO901317, and Orlistat, had no or only little effect on cell viability or the lipid phenotype (Figure 4 and Additional files 8, 9). This suggests that the T-47D cells are able to compensate the drug-induced lipid imbalance with several mechanisms, which would be compelling to study in depth.

## Conclusion

Taken together, our results highlight that tamoxifen resistant cell lines can potentially be used as a representative model for studies of tamoxifen-resistant patients. We propose that the breast cancer cells can acquire tamoxifen resistance by dysregulation of different cellular pathways, dependent on their individual molecular phenotypes. Here, we highlight the inhibition of lysosomal membrane permeabilization as one of the mechanisms to avoid cell death, whereas an increase in neutral lipids may enable the further survival of these cells. We further propose that drugs targeting cellular antioxidation machinery may be able to overcome tamoxifen resistance. However, investigating the relevance of the proposed mechanism of acquired resistance in patients remains a challenge. Given the vulnerability of tamoxifen resistant cells to approved drugs such as disulfiram and dasatinib, it would be interesting to investigate whether these compounds could also be effective in clinical trials in tamoxifen-resistant breast cancer patients.

## List of abbreviations

ER, estrogen receptor; DSRT, drug sensitivity and resistance testing; LMP, lysosomal membrane permeabilization, ROS, reactive oxygen species

## Acknowledgments

We acknowledge Prof. Marja Jäättelä (Danish Cancer Society Research Institute, Denmark) for comments to lysosomal membrane permeabilization assays. Anna Uro from the Ikonen laboratory (University of Helsinki) is acknowledged for technical assistance in biochemical lipid determination. High Throughput Biomedicine (HTB) unit (Laura Turunen, Maria Nurmi and Swapnil Podar), Sequencing unit (Pirkko Mattila), and High Content image Analysis (HCA) unit at the Institute for Molecular Medicine Finland (FIMM), HiLIFE, University of Helsinki, and Biocenter Finland are acknowledged of the high throughput drug profiling and high content imaging expertise. Piia Mikkonen is acknowledged for technical help with the drug screens.

## Authors’ contributions

SH and VP designed the study. SH performed experiments, data analysis and wrote the manuscript. MK performed RNA-Seq analysis and contributed to writing the manuscript. LP analyzed the lipid screen and contributed to writing the manuscript. RMK performed the LMP experiment. EI provided resources for the biochemical assay and participated in editing the manuscript. SK provided resources for RNA-sequencing and participated in editing the manuscript. VP and OK supervised the project and participated in manuscript writing and editing. All authors read and approved the final manuscript.

## Competing interests

The authors declare that they have no competing interests.

## Availability of data and materials

All data generated or analyzed during this study are included in this published article and its additional files. The raw and processed sequencing data have been deposited in the GEO database [GEO: GSE111151].

## Ethics approval and consent to participate

Not applicable.

## Consent for publication

Not applicable.

---

**Additional figure 1.**
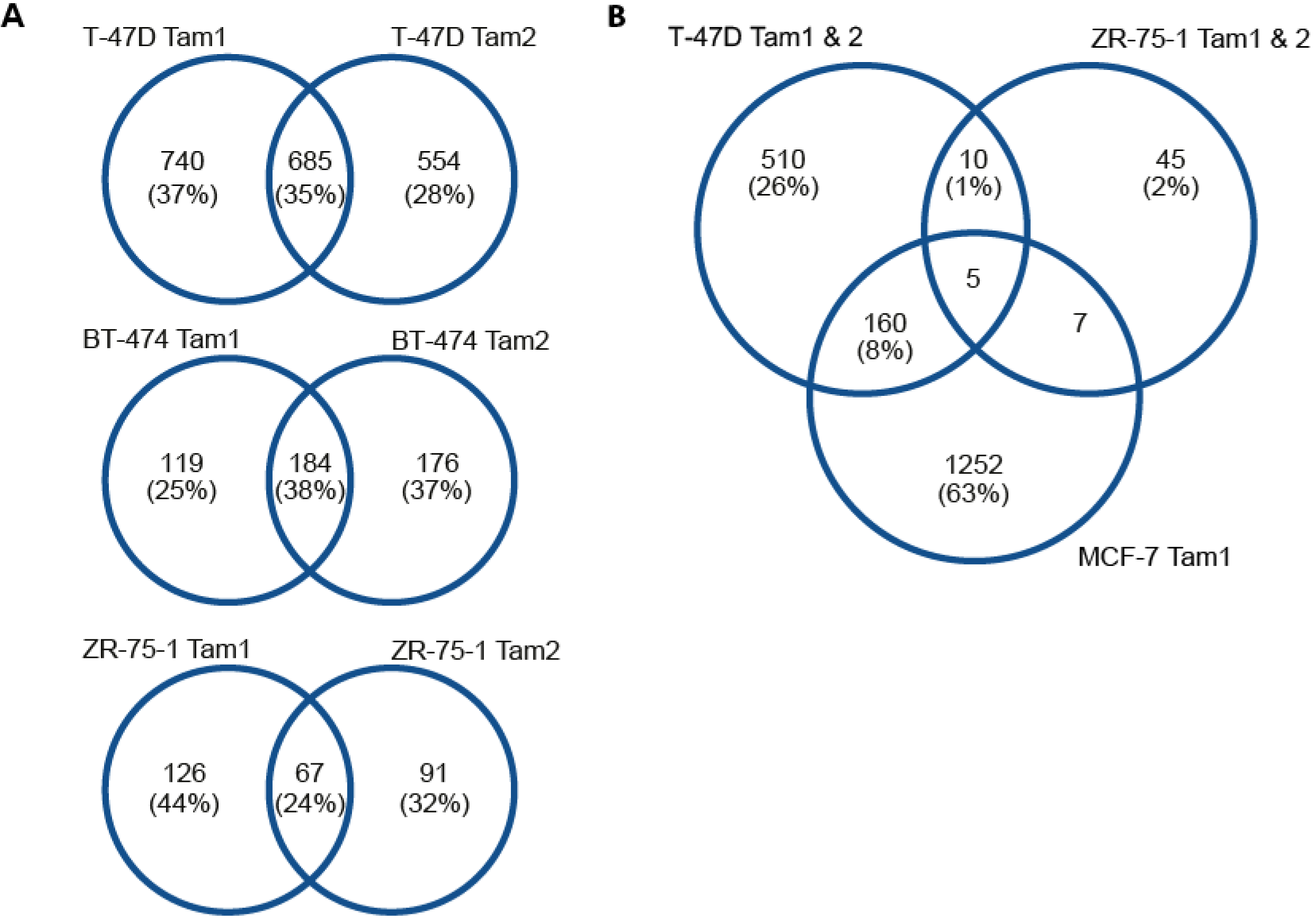
Tamoxifen-resistant cell lines display distinct expression changes. Tamoxifen-resistant clones derived from same parental cells (**A**) and of the luminal A subtype (**B**) differ in their expression changes. Venn diagrams show overlap in numbers and percentage of genes that are differentially expressed.

**Additional file 1**: Antibodies used in this study

**Additional file 2**: Drugs used in this study

**Additional file 3**: Statistical analysis of triglycerides, free cholesterol and cholesterol esters, LMP assay, and LipidToxGreen staining

**Additional file 4**: RNA-sequencing statistics

**Additional file 5**: Detected fusion genes of parental and tamoxifen-resistant cell lines

**Additional file 6**: Batch corrected CPM counts of cell lines and patients

**Additional file 7:** Enriched pathways with adjusted p<0.001

**Additional file 8**: DSS scores of CTG measurement and cell count

**Additional file 9:** Mean of the mean LipidToxGreen intensity values

